# The vascularised chamber device significantly enhances the survival of transplanted liver organoids

**DOI:** 10.1101/2023.04.24.538062

**Authors:** Denis D. Shi, Evelyn Makris, Yi-Wen Gerrand, Pu-Han Lo, George C. Yeoh, Wayne A. Morrison, Geraldine M. Mitchell, Kiryu K. Yap

## Abstract

Organoid transplantation has a promising future in the treatment of liver disease, but a major limitation is the lack of guidance on the most appropriate method for transplantation that maximises organoid survival. Human induced pluripotent stem cell (hiPSC)-derived liver progenitor cell organoids were transplanted into four different transplantation sites in a mouse model of liver disease, using five organoid delivery methods. Organoids were transplanted into the vascularised chamber device established in the groin, or into the liver, spleen, and subcutaneous fat. For organoid transplantations into the liver, organoids were delivered either in Matrigel alone, or in Matrigel and a polyurethane scaffold. At 2 weeks post-transplantation, the vascularised chamber had the highest organoid survival, which was 5.1x higher than the site with second highest survival (*p*=0.0002), being the intra-hepatic scaffold approach. No organoid survival was observed when delivered into the liver without a scaffold, or when injected into the spleen. Very low survival occurred in transplantations into subcutaneous fat. Animals with the vascularised chamber also had the highest levels of human albumin (0.33 ± 0.09 ng/mL). This study provides strong evidence supporting the use of the vascularised chamber for future liver organoid transplantation studies, including its translation into clinical therapy.

## INTRODUCTION

Recent pre-clinical advances in regenerative medicine suggest that organoid transplantation may alleviate liver disease [1, 2, 3]. However, transplantation approaches used in experimental animal models are often unsuitable for clinical translation because they are too invasive or risky, provide inadequate space for large-scale organoid delivery, or cannot be easily monitored and surgically explored. Examples of such methods include the cranial window [4, 5], dorsal skinfold chamber [6], and direct puncture or subcapsular delivery into organs such as the spleen [3], liver [7], or kidney [2]. To develop a clinically viable method for organoid transplantation, it is important to consider the surgical technique (level of difficulty and risk) and the location of organoid delivery (intra-hepatic or ectopic). Moreover, it is likely that different methods are more suited for different types of patients. For example, patients with early-stage disease may be adequately treated with a low-dose of organoids via a simple portal vein infusion, whereas patients with severe liver disease may require a greater number of organoids delivered via a more invasive method. To guide these future decisions in clinical therapy, the field requires experimental data comparing different methods and sites of organoid transplantation for liver disease. However only few such studies exist [4, 7, 8], with conflicting outcomes. There is a need for expanded studies addressing this pertinent issue.

In this study, human induced pluripotent stem cell (hiPSC)-derived liver organoids were transplanted into 4 different sites in a mouse model of liver disease. The sites were intra-hepatic onto the liver surface, spleen, subcutaneous fat, and the vascularised chamber created around the blood vessels in the groin. For intra-hepatic delivery, organoids were delivered either in a hydrogel plug smeared onto the surface, or a clinically used bioabsorbable porous scaffold [9, 10]. A unique aspect of the study is the inclusion of a vascularised chamber, which comprises a non-collapsible silicone chamber that acts as a protective device around a major artery and vein, inducing a highly angiogenic environment that robustly supports the engraftment of transplanted cells and tissues [11]. Although the vascularised chamber has been used to transplant various liver cell types [12, 13, 14, 15], its use has never been directly compared to other sites of transplantation, either for liver bioengineering or any other application.

## METHODS

The overall experimental workflow is shown in **Figure 1A**.

**Figure 1.**
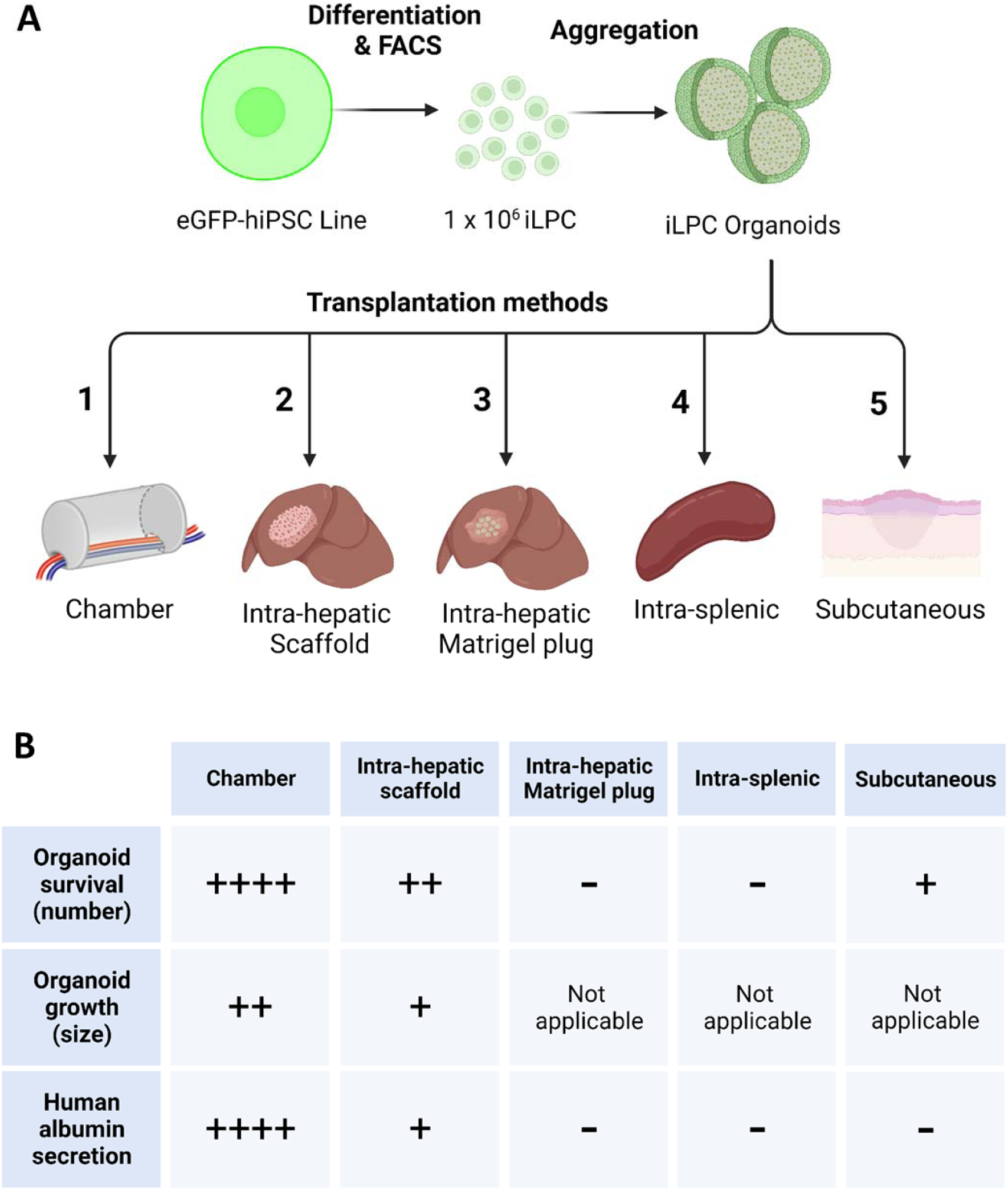
Study overview. **(A)** Schematic diagram of the experimental plan. **(B)** Summary of overall results.

### eGFP-hiPSC culture and differentiation into iLPCs

An eGFP reporter hiPSC line was used for this study, which has been previously characterised [16]. This eGFP-hiPSC line was generated by inserting the eGFP reporter gene into the GAPDH locus, thereby ensuring all cells constitutively express eGFP. hiPSCs were maintained in TeSR-E8 Media (STEMCELL Technologies, Vancouver, Canada) with 10% knockout serum replacement (ThermoFisher, Massachusetts, USA).

eGFP-hiPSCs were differentiated into hiPSC-liver progenitor cells (iLPCs) by plating onto Matrigel-coated plates (hESC-qualified Matrigel, Corning, New York, USA), then differentiated using a 12-day protocol with 3 steps. Base media throughout this period was RPMI-1640 media (ThermoFisher) with 1% B27 supplement (ThermoFisher). For the first 4 days, media was supplemented with 100ng/mL Activin A (Peprotech), then for the next 4 days media was supplemented with 50ng/mL bone morphogenetic protein 4 (BMP4) (Peprotech), and for the final 4 days 50ng/mL of hepatocyte growth factor (HGF) (Peprotech) was added. Following differentiation, cells were purified using fluorescence-activated cell sorting (FACS) on a BD Aria Fusion cell sorter (BD Bioscience, New Jersey, USA) using a GCTM-5 antibody conjugated to Allophycocyanin (APC) [GCTM-5 hybridoma provided by Professor Martin Pera (Jackson Laboratory) and Dr Alison Farley (Walter and Eliza Hall Institute)]. GCTM-5 has been previously shown to be expressed in endoderm/liver progenitor cells from adult liver as well as ESC-derived hepatoblasts [17, 18, 19, 20].

Purified GCTM-5+ iLPCs were expanded in LPC medium comprising base medium of Williams’ Medium E (ThermoFisher), with 10% foetal calf serum (Sigma-Aldrich, Missouri, USA), 10mM HEPES (ThermoFisher), 2mM GlutaMAX (ThermoFisher), 1% antibiotic-antimycotic (ThermoFisher), 5% R-spondin1 conditioned medium (supplied by Monash Biodiscovery Institute Organoid Program, Victoria, Australia), 10μg/mL Actrapid® human insulin (Novo Nordisk, Bagsværd, Denmark), 20ng/mL epidermal growth factor (EGF) (Peprotech), 30mg/mL insulin-like growth factor II (IGF-II) (Peprotech), 25ng/mL HGF (Peprotech), 50ng/mL fibroblast growth factor-10 (FGF) (Peprotech), 1.25mM N-acetyl cysteine (Sigma), 10mM Nicotinamide, 5µM A83-01 (Sigma), 10µM Forskolin (Abcam, Cambridge, United Kingdom), 10nM [Leu15]-Gastrin I (Sigma), and 0.5µM CHIR99021 (Sigma).

### Formation of iLPC organoids

iLPCs were lifted from monolayers, and 1 x 10 ^6^ iLPCs were plated onto each well of an ultra-low attachment 6-well plate (Corning). LPC medium for this step was modified to increase the R-Spondin1 concentration to 10%, with further additions of 20% Wnt3A conditioned medium (Monash), 25ng/mL Noggin (Peprotech), and 10µM of Y-27632 (Abcam). Individual cells were allowed to aggregate into organoids over a 48-hour period.

### Organoid processing for histology

iLPC organoids formed after 48 hours post-seeding were fixed for 3 hours in 4% paraformaldehyde (PFA) in phosphate buffered saline (PBS) (Sigma), before a double-embedding process in 1.5% agarose gel then paraffin as described previously [13, 14, 15]. Organoids were sectioned at 5µm thickness and mounted onto polysine-coated adhesion slides (Epredia, Michigan, USA).

### Transplantation of iLPC organoids

All animal experiments were performed with approval of the St Vincent’s Hospital Melbourne Animal Ethics Committee (Protocol AEC 20/19), in accordance with Australian National Health and Medical Research Council guidelines. *Fah*^−/−^*/Rag2*^−/−^*/Il2rg*^−/−^ (FRG) mice (Yecuris corporation, Oregon, USA) were maintained as described previously [15]. Animals were routinely maintained on 8mg/L of (2-(2-nitro-4-fluoromethylbenzoyl)-1,3-cyclohexanedione) (NTBC) supplemented in drinking water. From three weeks prior to transplantation and after organoid transplantation, NTBC was with-held every 5-7 days for a period of 2 days, and this cycle was continued until harvest, to induce mild liver injury and therefore to mimic the process of metabolic liver disease [15, 21].

Male and female mice (approximate 1:1 ratio) weighing 16-20g and 6-8 weeks of age were used for all experiments. A total of 5 different groups (N=3-7 per group) was completed, with each group representing a different type of organoid delivery and surgery. For all animals iLPC organoids generated from ^6^ 1cexlls w1 a0s delivered, suspended in 25µL of growth-factor full Matrigel (Corning).

For the vascularised chamber device group, chambers were established in the groin using a surgical method described previously [11, 13, 14, 15, 22]. Briefly, this involved surgically isolating the proximal segment of inferior epigastric pedicle (an artery/vein branch from the femoral vessels) by freeing it from its surrounding tissue, then sleeving the exposed portion with a 5mm length of split silicone tubing (Corning) and securing the tubing to the underlying muscle with 10-0 nylon suture (Ethicon, New Jersey, USA). The total chamber volume was 50µL. The proximal end and longitudinal seams of the tube were sealed using bone wax (Ethicon), then the chamber was filled from its distal end with 50µL of growth factor full Matrigel. The distal end was then sealed with bone wax to create an enclosed space. The skin was closed using staples. Chambers were allowed to vascularise through angiogenic sprouting from the enclosed artery/vein for a period of 3 weeks, then opened again for organoid transplantation. By this time some Matrigel had remodelled and was resorbed to provide enough space for 25µL of additional Matrigel containing iLPC organoids to be delivered.

For intra-hepatic organoid delivery in a scaffold and Matrigel, a polyurethane scaffold (NovoSorb® provided by PolyNovo Ltd, Port Melbourne, Australia) was used. This is a porous synthetic polymer containing interconnected pores of 300-600µm diameter. iLPC organoids suspended in 25µL of Matrigel were seeded into scaffold discs of 1mm thickness and 5mm diameter, and allowed to set at 37°C. For transplantation into the liver, a transverse laparotomy was performed, then an incision to the left liver lobe was made and a pocket was created with micro-forceps. A single scaffold was laid into this pocket and sandwiched between the left liver lobe and the overlying median lobe without sutures. This positioning meant that the sandwiched scaffold was unlikely to move from the pocket. The abdominal wall and skin were closed using a 7-0 prolene suture (Ethicon).

For intra-hepatic organoid delivery with Matrigel only, organoids in Matrigel were allowed to set at 37°C to form a semi-solid plug. A pocket was created in the left liver lobe as described above, and the Matrigel plug containing organoids was smeared into this pocket using a spatula. Cauterisation was used to leave a small scar on the edge of the lobe so the pocket could be identified during harvest.

For subcutaneous organoid delivery, organoids in Matrigel were injected into the subcutaneous fat pad in the groin using a 29G insulin syringe. This created a bleb just under the skin, confirming subcutaneous delivery (and not muscle delivery). The needle insertion point was sutured with a 7-0 prolene suture to prevent organoid leakage.

For intra-splenic organoid delivery, an incision was made into the left flank, and the spleen was partially exposed to access the distal pole. Organoids suspended in Matrigel were injected into the spleen using a 29G insulin syringe inserted from the distal pole, and the tip of the spleen proximal to the needle entry-point was tied off using a 5-0 vicryl suture (Ethicon) to prevent organoid leakage. The abdominal wall and skin were closed using a 7-0 prolene suture.

### Post-transplantation tissue harvest

Tissues were collected 2 weeks post-transplantation and fixed in 4% PFA in PBS (Sigma) overnight prior to embedding in paraffin. Tissue sections of 5µm thickness were cut and mounted onto polysine adhesion slides (Epredia, Michigan, USA).

### Immunohistochemistry

Paraffin sections of *in vitro* organoids and *in vivo* tissue samples were dewaxed and rehydrated, then immuno-labelled using a standard protocol. This involved sequential treatment with heat-mediated citric acid (Sigma) buffer antigen retrieval, peroxidase quench with 3% hydrogen peroxide (Merck, New Jersey, USA), protein block with Dako protein block solution (Dako, Glostrup, Denmark), incubation with primary antibodies [**eGFP** (Abcam, ab6673, 1:400), **HNF4α** (R&D Systems, Minnesota, USA, PP-H14150C, 1:200), **Sox9** (Abcam, ab185966, 1:1000), **human albumin** (Bethyl Laboratories, Texas, USA, A80-229A, 1:200), **K i 6** (**7**ThermoFisher, RM-9106-S1, 1:300), **CD31** (ThermoFisher, PA1-37326, 1:50) or **LYVE-1** (Abcam, ab14917, 1:250)] in Dako antibody solution (Dako), incubation with secondary antibodies (either anti-goat, anti-rabbit, or anti-mouse biotinylated secondary antibody, all from Vector Labs, California, USA, at 1:200), detection using a Vector Elite ABC kit (Vector), then 3,3’-Diaminobenzidine (DAB) chromogen staining (Dako). Sections were counter-stained with haematoxylin and cover-slipped with Entellan mounting medium (Merck).

### Morphometry assessments

The number and size of eGFP+ liver organoids per tissue sample, and the percentage of HNF4α+ and Sox9+ nuclei within organoid structures were identified using serial paraffin sections immuno-labelled with eGFP, HNF4α and Sox9. For each tissue sample, serial sections were first staine regular intervals (every 10 sections) with haematoxylin & eosin to identify the area of greatest engraftment. Tissue sections from this area were then stained with the relevant markers for morphometric analysis.

Organoids were identified using 4x magnification, then individually imaged at 40x magnification for assessment of organoid diameter, circumference, and area, and counting of HNF4α positive/negative and Sox9 positive/negative nuclei within organoids. Number of organoids per sample and nuclei counts were conducted manually, whereas ImageJ software (National Institutes of Health, Maryland, USA) was used to assess organoid diameter, circumference, and area. Morphometric assessments were completed by one observer, then independently checked by a second observer. Between observers there was less than 5% of discrepancy, so the first observer’s assessments have been used for analysis.

### Human albumin ELISA

During the harvest of tissues post-transplantation, mouse blood was collected via intra-cardiac puncture into lithium heparin separator tubes (Greiner Bio-one, Kremsmünster, Austria) and centrifuged to obtain plasma. Mouse plasma was analysed for human albumin using a commercial ELISA kit (Bethyl Laboratories, Texas, USA), following manufacturer’s instructions with the inclusion of positive and negative controls.

### Statistical analysis

Data is presented as mean ± SEM and analysed with one-way ANOVA and Bonferroni post-hoc analysis using GraphPad Prism v9.0 (GraphPad Software, California, USA). N=3-7 was included for all groups.

### Data availability statement

The data that support the findings of this study are available from the corresponding author upon reasonable request.

## RESULTS

### iLPCs can be aggregated into organoids that display key liver markers in vitro

eGFP+ GCTM-5+ iLPCs aggregated into compact semi-spherical organoids within 48 hours of plating into ultra-low adhesion plates. Organoids ranged in diameter from 50-150µm, with the majority being approximately 100µm **(Figures 2A, 2B**. W) hen organoids were fully formed at 48 hours, most cells within organoids were positive for the two major nuclear transcription factors regulating hepatic function (HNF4α) **(Figure 2C)** and biliary differentiation (Sox9) **(Figure 2D)**. Cells were tightly packed within organoids, although small cystic spaces often occurred throughout organoids **(Figures 2C-2F).** Albumin was only found in a small number of cells within organoids **(Figure 2E)**, and over 50% of cells were positive for the proliferation marker Ki67 **(Figure 2F)**. Organoids could be resuspended in Matrigel and seeded into circular polyurethane scaffold discs **(Figure 2G)**, and eGFP+ organoids were confirmed to be dispersed throughout the pores of the scaffold **(Figure 2H).**

**Figure 2.**
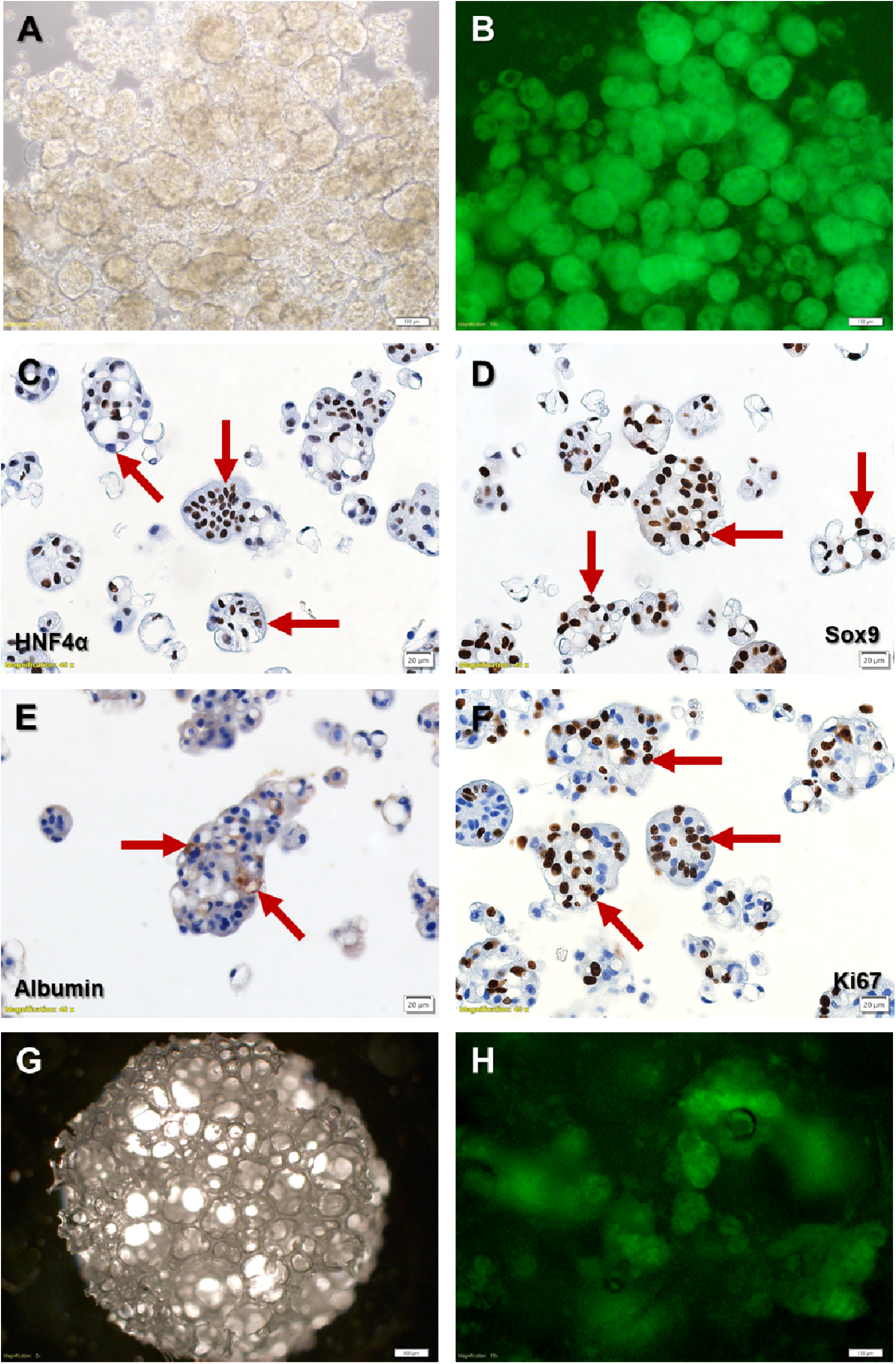
*In vitro* iLPC organoids. **(A)** Bright-field imaging of iLPC organoids at day 48 post-cell seeding. **(B)** Corresponding fluorescence imaging of the same organoids showing eGFP expression in all organoids. **(C)** HNF4α immunohistochemistry, showing robust HNF4α expression in most nuclei (red arrows). **(D)** Sox9 immunohistochemistry, showing robust Sox9 expression in most nuclei (red arrows). **(E)** Albumin immunohistochemistry, showing low expression in only a small number of cells within organoids (red arrows). **(F)** Ki67 immunohistochemistry, showing that >50% of cells were Ki67+ (red arrows) and proliferative. **(G)** The NovoSorb® polyurethane scaffold used for intra-hepatic organoid delivery, which were circular discs 5mm in diameter, 1mm in thickness, with interconnected pores 300-600µm in diameter. **(H)** eGFP+ iLPC organoids were suspended in Matrigel and seeded into the polyurethane scaffold and were found well dispersed throughout the pores of the scaffold. **Scalebars** -20µm for **(C)**, **(D)**, **(E)**, **(F)**, 100µm for **(A)**, **(B)**, **(H)**, 500µm for **(G)**.

Collectively, a relatively uniform collection of organoids was generated from 1 x 10 ^6^ iLPCs seeded into a single well, and these structures contained cells that displayed key liver marker s Sox9, albumin). Co-expression of HNF4α and Sox9 indicated relative immaturity and the bipotential capacity of iLPCs to differentiate into both hepatocytes and cholangiocytes, and the high expression of Ki67 implies that the organoids have substantial capacity for expansion *in vitro* and were proliferative at the time of *in vivo* transplantation.

### Extensive engraftment of hepatobiliary structures occurs within the vascularised chamber

Morphometric analysis and functional analysis (human albumin ELISA) for all sites is presented collectively at the end of the results section, after a description of each of the transplantation sites.

Organoids were transplanted into the vascularised chamber 3 weeks after initial establishment, where a small amount of vascularised tissue and remnant Matrigel surrounded the enclosed pedicle with enough space to introduce organoids suspended in 25µL of new Matrigel. At 2 weeks post-transplantation of organoids, cylindrical-shaped tissue moulded by the confines of the silicone chamber tubing was harvested, which contained large areas of moderately cellular remnant Matrigel infiltrated with adipocytes, inflammatory cells, and small blood vessels surrounded the main pedicle **(Figure 3A).** All tissue constructs were encapsulated by a densely cellular, vascularised connective tissue layer **(Figure 3A)**. Adjacent to the Matrigel and fat area, many organoids were clustered together, easily identified as loosely circular and oval structures with cuboidal-columnar epithelium lining large dilated cystic spaces containing heterogeneous acellular secretions **(Figure 3A, 3B)**.

**Figure 3.**
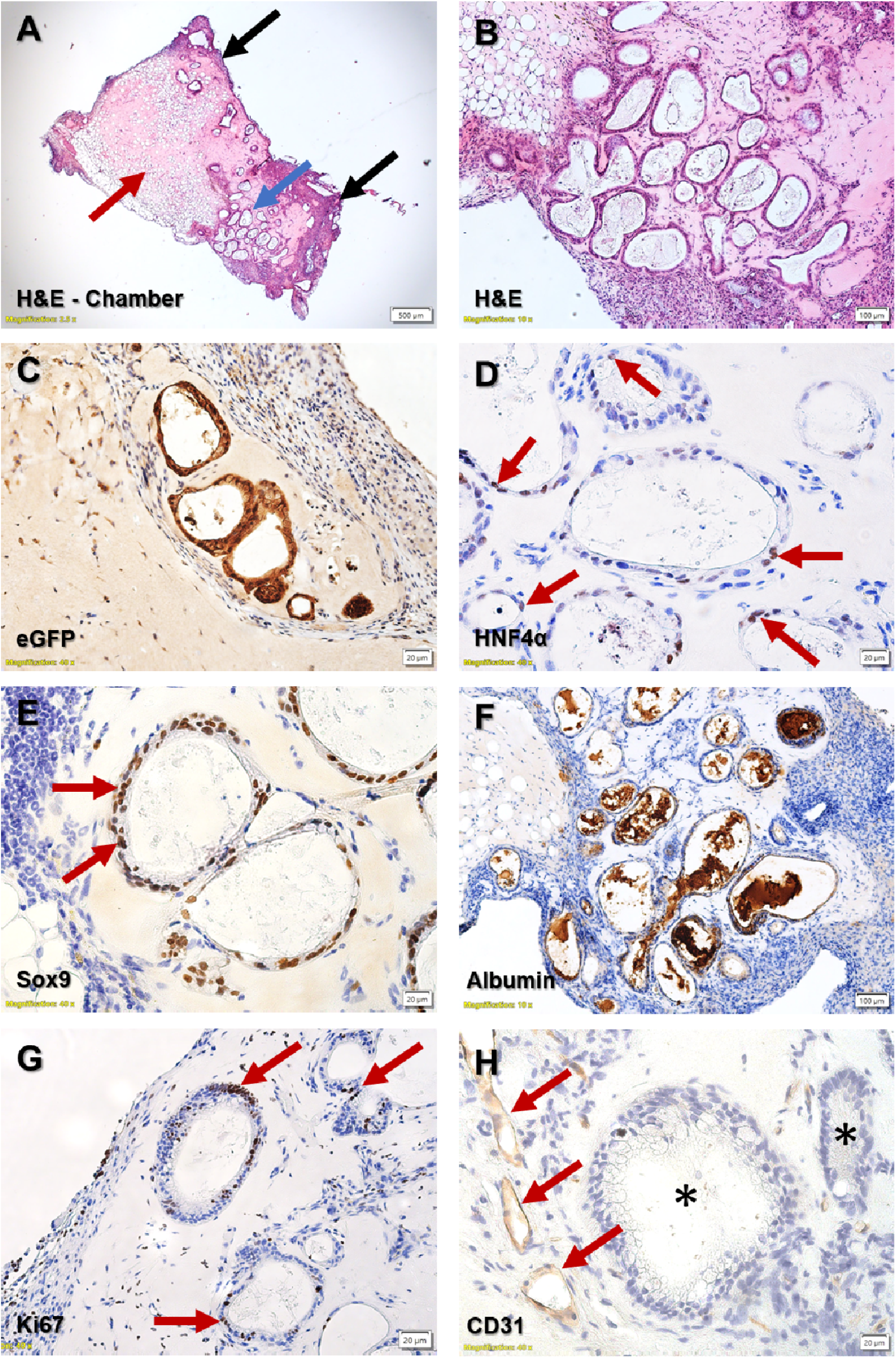
Organoid transplantations into the vascularised chamber. **(A)** Low magnification haematoxylin & eosin (H&E) staining of the chamber tissue, demonstrating it was encapsulated by a layer of inflammatory connective tissue (black arrows), had large areas of sparsely cellular remnant Matrigel infiltrated with native adipocytes (red arrow), and areas where multiple organoids were clustered together (blue arrow). **(B)** High magnification H&E staining of thechambe demonstrating multiple organoids which formed ductular structures which contained large cystic spaces with heterogeneous secretions lined by cuboidal-columnar epithelium. **(C)** Organoids were identified by their expression of eGFP. **(D)** Only approximately 50% of cells within organoids were HNF4α+. **(E)** The majority of cells within organoids were Sox9+. **(F)** The heterogeneous intra-luminal secretions within organoids labelled strongly for human albumin. **(G)** Up to 40% of cells within organoids were Ki67+ and proliferative. **(H)** Organoids were often near CD31+ native blood vessels in the chamber. **Scalebars** -20µm for **(C)**, **(D)**, **(E)**, **(G)**, **(H)**, 100µm for **(B)**, **(F)**, 500µm for **(A)**.

Organoids were found in every sample of the group (N=3). Although different in appearance to the more tightly packed circular organoids *in vitro*, these structures were confirmed to be the xenografted iLPC organoids due to their eGFP expression **(Figure 3C).** In contrast to *in vitro* organoids, HNF4α expression was found in a much lower proportion of cells **(Figure 3D)**, although Sox9 expression was still found in most cells **(Figure 3E)**. The large cystic secretions were strongly positive for albumin **(Figure 3F)**, and a substantial number of cells within the organoids remained Ki67+ (**Figure 3G).** Additionally, CD31+ blood vessels were often found near organoids **(Figure 3H).**

These results indicate the robust engraftment of organoids in the chamber. Organoids formed proliferative and dilated hepatobiliary structures which co-expressed hepatocyte and biliary markers, secreted copious amounts of intra-luminal albumin, and interacted with native blood vessels developing within the chamber. The striking change in morphology and increase in albumin expression suggests developmental progression compared to *in vitro* organoids.

### Intra-hepatic delivery in a scaffold leads to enhanced organoid engraftment

Two methods of organoid delivery onto the surface of the liver were tested. Organoids were either suspended in Matrigel and seeded into a polyurethane scaffold which was applied onto the liver surface and sandwiched between two liver lobes or smeared onto the liver surface as a semi-solid plug of Matrigel without the use of a scaffold.

When the scaffold was used, it was found tightly sandwiched between the liver lobe it was applied onto and the overlying liver lobe. Histology demonstrated that the scaffold was well adhered to both liver lobes, with native tissue (mostly inflammatory cells and blood vessels) infiltrating the scaffold pores **(Figure 4A, 4B)**. Only a few organoids per sample were observed, and organoids were present in 4 out of 6 samples (N=6). Organoids were confirmed to be eGFP+ **(Figure 4C)**, and similar to organoids in the chamber co-expressed HNF4α **(Figure 4D)** and Sox9 **(Figure 4E)**, secreted intra-luminal albumin **(Figure 4F)**, and contained Ki67+ proliferative cells **(Figure 4G).** Where organoids were present, they were often found in the periphery of scaffolds close to the native mouse liver tissue with LYVE-1+ blood vessels extending from the liver into the scaffold area to surround the organoids **(Figure 4H)**. LYVE-1 was used to identify blood vessels because liver sinusoidal endothelial cells (LSECs) which line liver-specific blood vessels can be identified with this marker.

**Figure 4.**
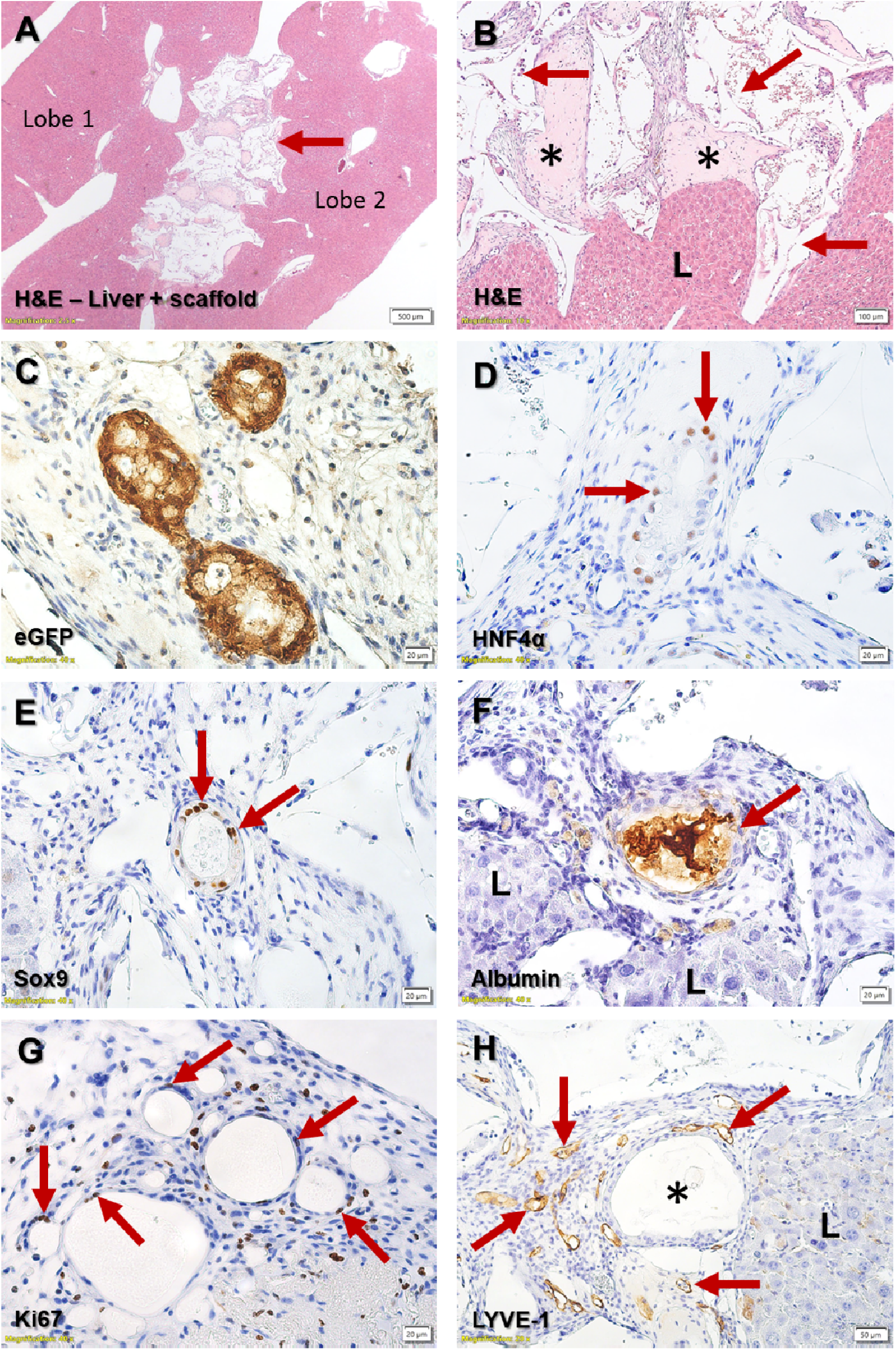
Organoid transplantations into the liver using a polyurethane scaffold. **(A)** Low magnification H&E staining demonstrating the scaffold (red arrow) well adhered between two liver lobes. **(B)** High magnification H&E demonstrating the scaffold directly adjacent and tightly adhered to the native mouse liver (labelled L), with Matrigel, infiltrating host tissue within the pores of the scaffold (labelled with asterisks), and the walls of the scaffold pores appearing as clear triangular material throughout the scaffold area (red arrows). **(C)** eGFP+ organoids were identified in the scaffold. **(D)** Many of the cells in the organoid were HNF4α (red arrows). **(E)** Most of the cells in the organoid were Sox9+ (red arrows). **(F)** The intraluminal secretions within organoids were strongly positive for human albumin (red arrow), and organoids were often found in the periphery of the scaffold close to native mouse liver (labe l(l**G**e)dSoLm). e of the cells within organoids were Ki67+ and proliferative (red arrows). **(H)** Organoids in the scaffold (asterisk) were often located near the native mouse liver (labelled L), and LYVE-1+ blood vessels from the liver extended into the scaffold to surround organoids (red arrows). **Scalebars** - 20µm for **(C)**, **(D)**, **(E)**, **(F)**, **(G)**, 50µm for **(H)**, 100µm for **(B)**, 500µm for **(A)**.

In contrast, when organoids were delivered in Matrigel alone, no organoids were found in any sample (N=7). The remnant Matrigel could be identified as a distinct and sparsely cellular layer on the liver surface, with a densely cellular inflammatory layer of cells overlying the Matrig e varied in thickness **(Figure 5A-5D)**.

**Figure 5.**
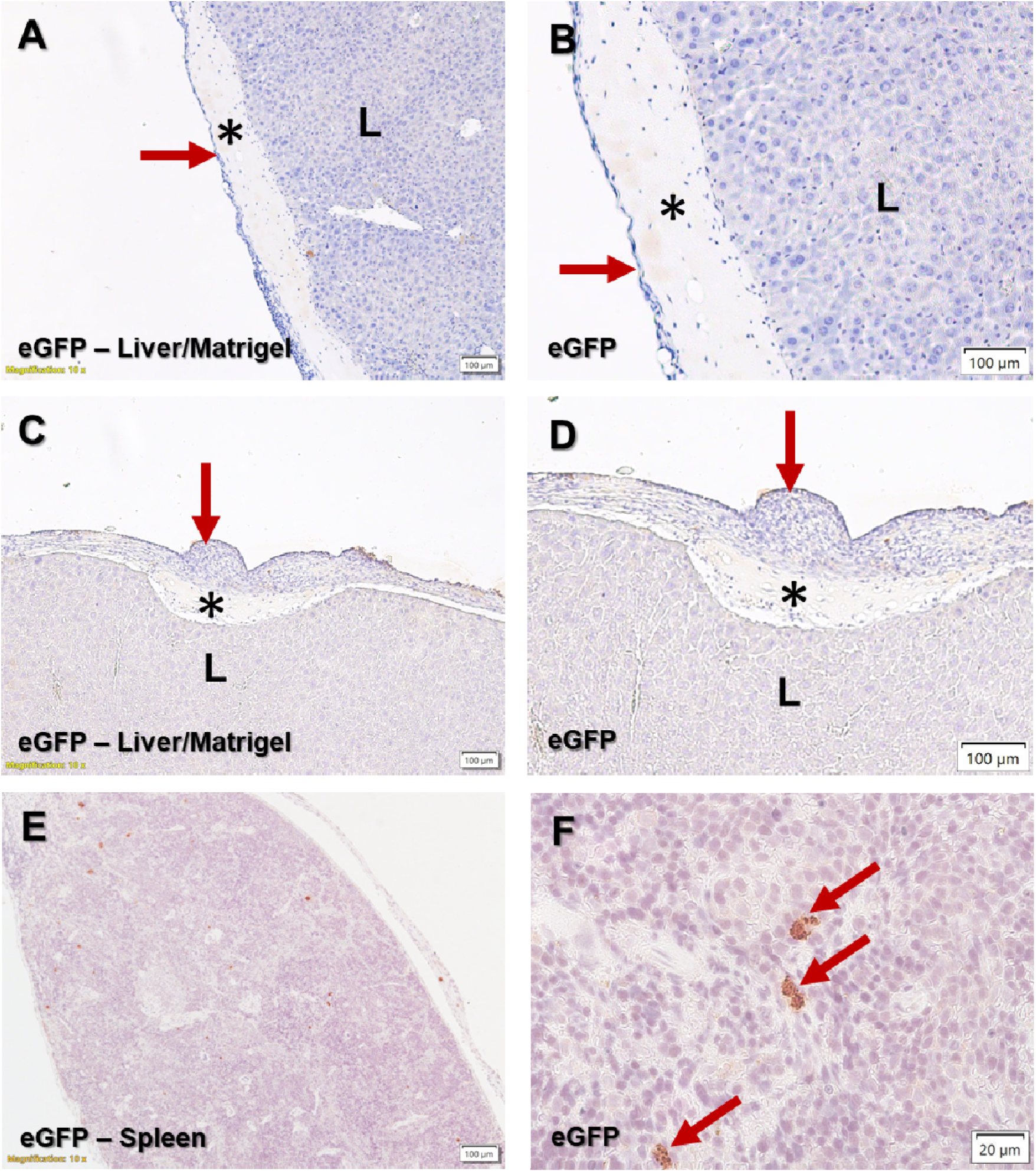
Organoid transplantations into the liver and spleen using Matrigel alone. **(A)** eGFP immunohistochemistry of livers transplanted with organoids in Matrigel alone resulted in no organoid survival. The remnant Matrigel was present on the surface of the liver (labelled L) as a thin sparsely cellular layer (asterisk), with a densely cellular inflammatory layer of cells overlaying the Matrigel (red arrow). **(B)** Higher magnification of eGFP immunohistochemistry. **(C)** In some livers the superficial layer of inflammatory cells was much thicker (red arrow), but the same architecture was present, which was a layer of remnant Matrigel (asterisk) on top of the host mouse liver (L), capped by a superficial layer of inflammatory cells (red arrow). **(D)** Higher magnification of eGFP immunohistochemistry in a liver with a thick layer of inflammatory cells on top of the remnant Matrigel. **(E)** In spleens transplanted with organoids, no organoid structures were present, demonstrated with eGFP immunohistochemistry. **(F)** There were isolated flecks of eGFP+ staining scattered throughout the spleen (red arrows), but this was not associated with nucleated cells. **Scalebars** - 20µm for **(F)**, 100µm for **(A)**, **(B)**, **(C)**, **(D)**, **(E)**.

Overall, organoid survival was diminished with intra-hepatic delivery, and engraftment was only found when organoids were delivered in a scaffold. This suggests that given the same site (liver surface) and extracellular matrix (Matrigel) and organoid dosage (total of 1 x 10 ^6^ cells), the use of the scaffold constrained organoids to a localised site and promoted their survival. The scaffold material was well adhered onto the liver at 2 weeks and showed dense infiltration of host tissue indicating that the material was being integrated over time.

### Intra-splenic organoid delivery leads to zero organoid engraftment

When organoids were suspended in Matrigel and injected into the spleen, no survival was found (N=4). Small, isolated specks of eGFP+ staining was scattered throughout the spleen **(Figure 5E, 5F)**, but were not associated with nucleated cells and this could either be remnants of disintegrated iLPCs or a staining artifact. Because cells injected into the spleen are commonly known to rapidly translocate to the liver by travelling through the vasculature and blood flow which drains from the spleen into the liver, the livers of animals that received intra-splenic organoid injections were systematically sampled. This involved the dissection of every lobe into 5mm strips (approximately 3-4 strips per lobe) and histological examination of every strip. This step confirmed no eGFP+ organoids were found in any animals, in both the spleen and liver. Moreover, liver translocation in this situation was not expected because organoids typically 50-150µm in diameter were unlikely to traverse into the splenic vasculature because of their size, and the use of viscous Matrigel which solidifies at body temperature would be a further impediment.

### Subcutaneous delivery leads to minimal organoid engraftment

During harvest, subcutaneous plugs of Matrigel could be identified as a pearly semi-translucent globule embedded in the subcutaneous fat just under the skin where a prolene marking suture had been used to close the needle puncture site during initial injection. With haematoxylin & eosin staining, the remnant Matrigel appeared as a pinkish often sparsely cellular area with infiltrating inflammatory cells, blood vessels, and fat, similar to the appearance in the vascularised chamber **(Figure 6A, 6G)**. Matrigel areas were encapsulated by subcutaneous fat and connective tissue **(Figure 6A, 6B)**, reflecting the en bloc resection technique used to harvest the tissue which included the Matrigel plug and a large margin of surrounding tissue. Amongst animals that received subcutaneous transplants (N=6), organoid survival was present in only 2 animals. In both animals only one organoid was observed, with very different characteristics. In the first animal, the eGFP+ organoid was very small (diameter 50.36µm, circumference 152.21µm, cross-sectional area 1658.33µm ^2^) **(Figure 6B)**, with only a moderate amount of intra-luminal albumin (denoted by staining intensity compared to organoids found in subcutaneous and chamber transplants) **(Figure 6C).** All cells in the organoids co-expressed HNF4α **(Figure 6D)** and Sox9 **(Figure 6E)**, with approximately 40% of cells being Ki67+ and proliferative **(Figure 6F)**. In contrast, in the second animal the organoid was 5x larger by diameter and circumference and 23x larger by area (diameter 251.85µm, circumference 726.75µm, cross-sectional area 38433.31µm ^2^) **(Figure 6H).** The organoid contained a larger area of intra-luminal albumin with slightly increased staining intensity **(Figure 6I)**. Importantly, the organoid completely lacked the expression of HNF4α **(Figure 6J)** and Sox9 **(Figure 6K)**, and only approximately 20% of cells were Ki67+ **(Figure 6L)**.

**Figure 6.**
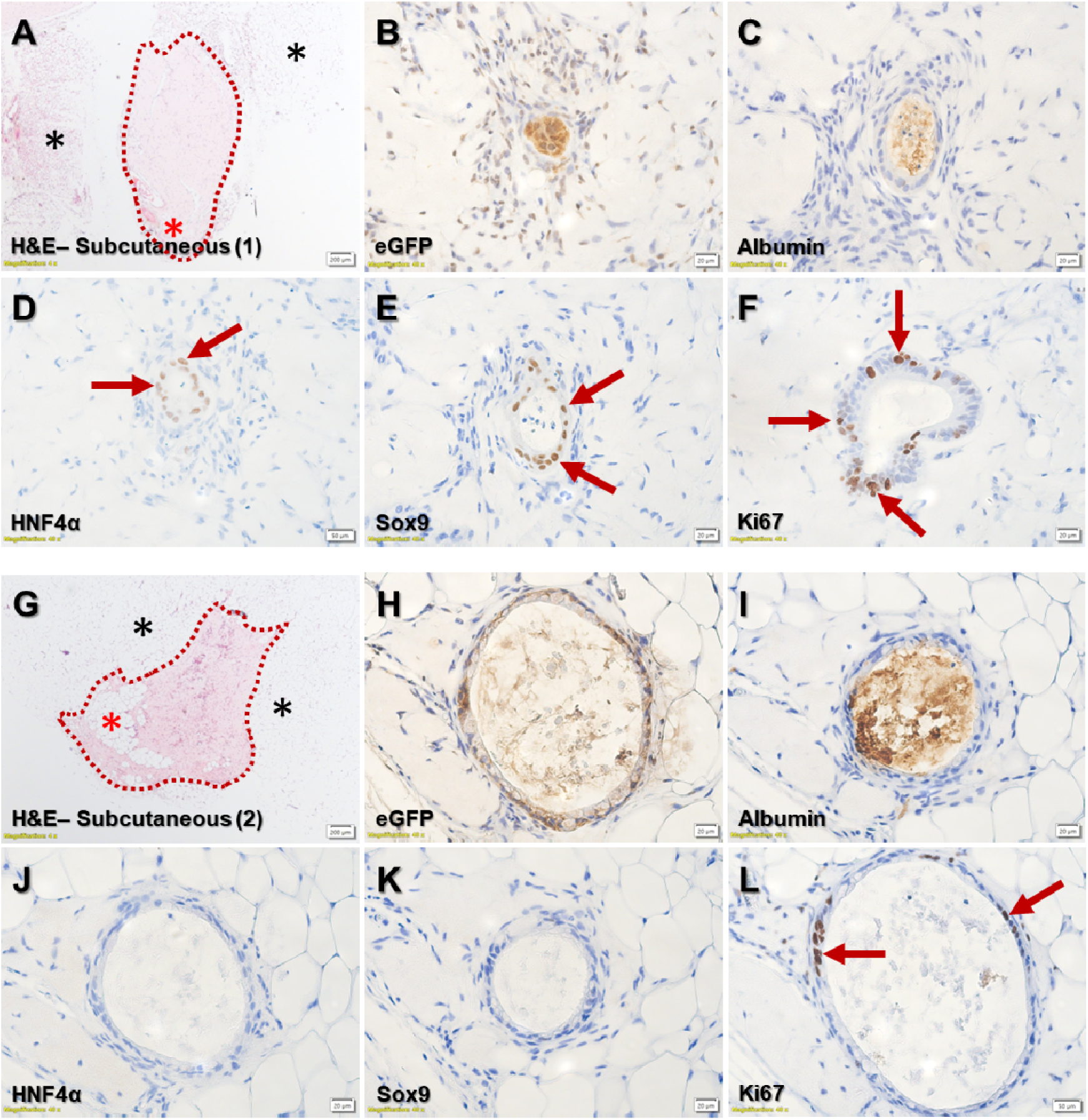
Organoid transplantations into subcutaneous fat. **(A)** H&E staining of the resected tissue demonstrating the remnant Matrigel seen as a pink sparsely cellular area (red dotted line) with some infiltrating adipocytes (red asterisk), clearly demarcated from the surrounding adipose and connective tissue (asterisks). Organoids were found in only 2 animals (out of 6), with each sample only containing 1 organoid. The 1^st^ sample is shown here. **(B)** In sample 1, only one small eGFP+ organoid was present. **(C)** Organoids contained intra-luminal human albumin. **(D)** All cells in the organoid were HNF4α+ (red arrows). **(E)** All cells in the organoid were Sox9+ (red arrows). **(F)** Approximately 20% of cells were Ki67+ and proliferative (red arrows). **(G)** H&E staining of sample 2, showing remnant Matrigel (dotted red line) with areas of infiltrating adipocytes (red asterisk), surrounded by subcutaneous adipose and connective tissue (black asterisks). **(H)** The eGFP+ organoid in this sample was much larger. **(I)** Similar to sample 1 the organoid in sample 2 also contained intra-luminal albumin. However, in contrast to sample 1 this organoid had no HNF4α+cells **(J)** or Sox9+ cells **(K)**. **(L)** Only a small proportion of cells were Ki67+. **Scalebars** - 20µm for **(B)**, **(C)**, **(E)**, **(F)**, **(H)**, **(I)**, **(J)**, **(K)**, 50µm for **(D)**, **(L)**, 200µm for **(A)**, **(G)**.

To summarise, subcutaneous transplantation led to very low engraftment of organoids with considerable variation between the organoids found in the two animals with engraftment.

### The vascularised chamber enhances organoid survival and growth compared to other sites

Comparing organoid survival between transplantation sites, one-way ANOVA indicated significant overall difference between transplantation sites (*p*<0.0001). The vascularised chamber had the highest number of eGFP+ organoids (24.67 ± 8.41 organoids) which was significantly higher than all other transplantation sites. The chamber had 5.1x the number of organoids than intra-hepatic delivery using a scaffold (4.83 ± 2.56 organoids, *p* = 0.0002), 74.8x compared to subcutaneous delivery (0.33 ± 0.21 organoids, *p*<0.0001), and was also higher than intra-hepatic delivery using Matrigel alone and intra-splenic delivery which had zero organoid survival in all animals in both groups (0 ± 0 organoids, *p*<0.0001) **(Figure 7A)**.

**Figure 7.**
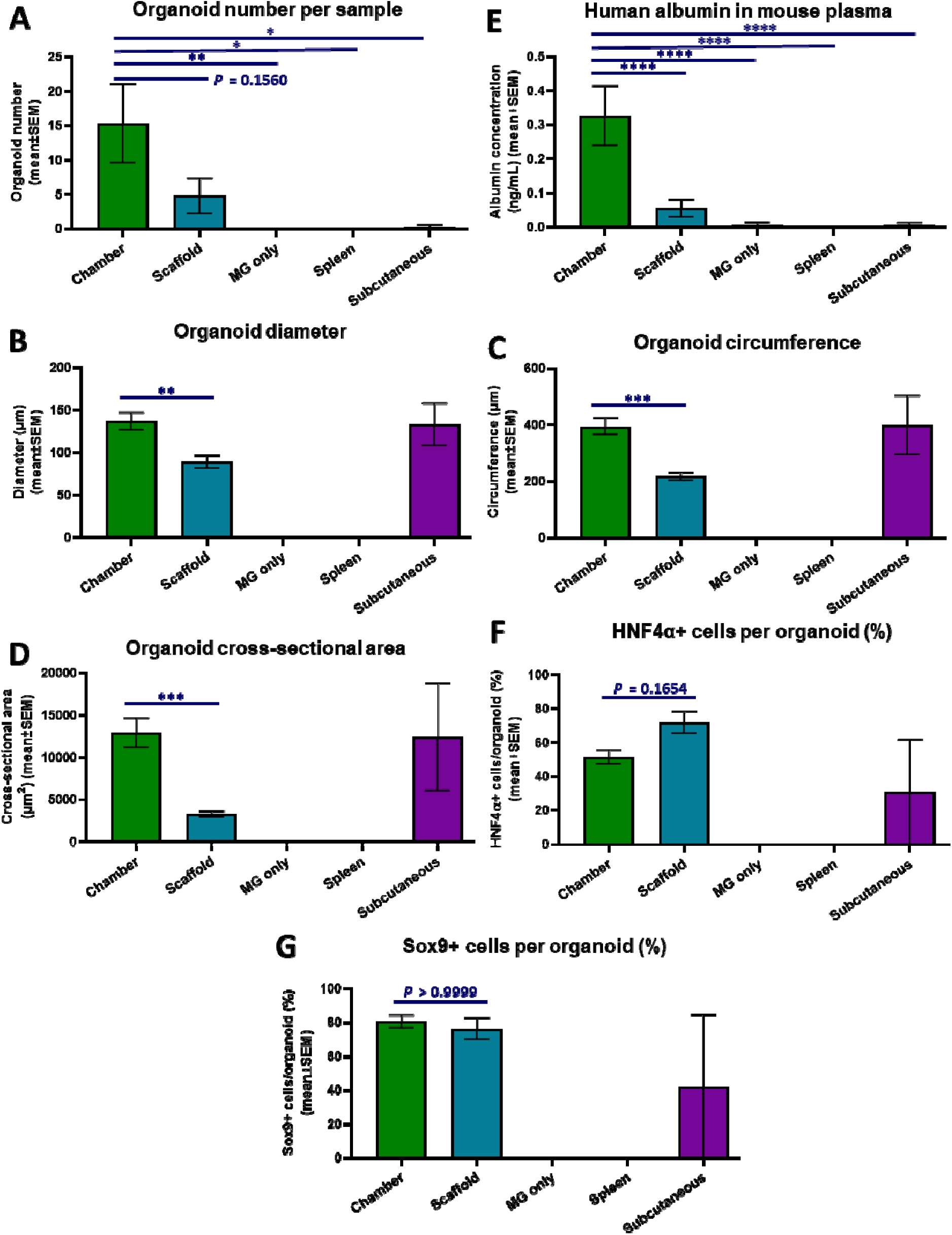
Morphometric & functional analysis. **(A)** Organoid survival (number of organoids per sample) **(B)** Human albumin levels in mouse plasma. **(C)**Organoid diameter. **(D)** Organoid circumference. **(E)**Organoid cross-sectional area. **(F)**Percentage of HNF4α+ cells per organoid. **(G)** Percentage of Sox9+ cells per organoid. Analysed with one-way ANOVA with Bonferroni post-hoc analysis. * *p*<0.05, ** *p*<0.005, *** *p*<0.0005, **** *p*<0.0001. N=3-7 per group for **(A)**, **(B)**, **(C)**, **(D)**, **(E)**, N=9-39 per group for **(F)**, and N=16-36 for **(G).**

This finding corroborated the levels of human albumin detected in mouse plasma, which was again found to have significant overall difference between transplantation sites (*p*<0.0001) and was significantly higher in the vascular is edchamber (0. 3 3 ± 0. 0 9 n g / m L transplantation sites. Animals with the chamber had 6x the amount of human albumin compared to intra-hepatic delivery using a scaffold (0.055 ± 0.03 ng/mL, *p*<0.0001), 47.8x compared to intra-hepatic delivery using Matrigel alone (0.0069 ± 0.007 ng/mL, *p*<0.0001), 49.3x compared to subcutaneous delivery (0.0067 ± 0.006 ng/mL, *p*<0.0001), and was also higher than intra-splenic delivery (0 ± 0 ng/mL, *p*<0.0001) **(Figure 7B)**.

The diameter, circumference, and cross-sectional area of organoids in the vascularised chamber were compared to organoids in the scaffold in the liver. A total of 53 organoids in the chamber group (from 3 animals), and 29 organoids in the intra-hepatic scaffold group (from 6 animals) were assessed for this analysis. Given the poor organoid survival in the subcutaneous delivery group and the marked variation between the two organoids assessed (from two different animals), this data was included in the graph but not considered meaningful for comparison or comment. With all three measurements of organoid size, organoids in the chamber were significantly larger than those in the intra-hepatic scaffold. Organoids were 1.5x larger in diameter (137.00 ± 10.07µm for chamber vs 89.16 ± 7.37µm for scaffold, *p* = 0.014) **(Figure 7C)**, 1.8x larger in circumference (394.6 ± 28.64µm for chamber vs 218.0 ± 13.69µm for scaffold, *p* = 0.0002) **(Figure 7D)**, and 4x larger in cross-sectional area (12967 ± 1722µm^2^ for chamber vs 3267 ± 365µm ^2^ for scaffold, *p* = 0.0006) **(Figure 7E)**.

Taken together, organoid transplantation into the vascularised chamber significantly increased survival (organoid number), function (human albumin ELISA), and individual organoid growth in terms of size (diameter, circumference, cross-sectional area) compared to all other transplantation methods and sites. The only other site/method with some organoid survival and function was intra-hepatic delivery with a scaffold, but this was very modest compared to the chamber.

### In vivo organoids in the chamber and intra-hepatic scaffold are still developmentally immature

Individual organoids in the chamber and scaffold groups were assessed for the proportion of HNF4α+ and Sox9+ cells to determine their level of maturity. Between the two groups a total of 48 organoids were assessed for HNF4α expression (N=9-39 per group), and 52 organoids for Sox9 (N=16-36 per group). For reasons mentioned previously, organoids in the subcutaneous group were included in the assessment but were not included in the overall interpretation of results.

With HNF4α, 51.53 ± 4.1% of cells in organoids were HNF4α+ in the chamber group, and 71.99 ± 6.4% of cells in organoids were HNF4α+ in the intra-hepatic scaffold group **(Figure 7F)**. Although not statistically significant (*p* = 0.1654), the 40% increase in HNF4α expression in the intra-hepatic scaffold group suggests that the native liver provides a hepato-inductive niche that is not present in the ectopic vascularised chamber.

With Sox9, 80.57 ± 3.8% of cells in organoids were Sox9+ in the chamber group, and 76.23 ± 6.3% of cells in organoids were Sox9+ in the intra-hepatic scaffold group (*p*>0.9999) **(Figure 7G)**. The similarly high expression of Sox9 in both groups (with co-expression of HNF4α) suggests that most cells in organoids *in vivo* remained at a liver progenitor cell stage. Upon terminal differentiation into mature hepatocytes, liver progenitor cells cease to express Sox9 and only express HNF4α as they have developed along the hepatic lineage.

## DISCUSSION

By comparing the survival and function of iLPC organoids transplanted using 5 different methods across 4 transplantation sites *in vivo*, this study demonstrated that the vascularised chamber led to the highest survival of organoids. Transplantation into the liver surface using Matrigel alone led to no survival, but when organoids were delivered with Matrigel into a scaffold, this led to modest survival although still 5.1x less than the chamber. Subcutaneous transplantation had minimal survival, and intra-splenic transplantation had zero survival. Human albumin levels in mouse plasma correlated with the survival of organoids, with the highest levels of human albumin detected in animals with the vascularised chamber. The results are summarised in the table presented in **Figure 1B**.

Previous studies compared 2-3 different transplantation sites, with conflicting data on the best site for organoid engraftment [4, 8], and only one study examined engraftment into the liver itself [7]. Including the liver site for comparison is important because delivering organoids to the native organ serves as a useful reference point. Furthermore, studies with cell transplantation have indicated that intra-portal infusion of cells into the liver leads to the best survival compared to other sites [23], although this specific method may not be suitable for organoids because of their size and thus increased risk of portal hypertension or portal vein thrombosis [24, 25]. In this study, two different methods were used to deliver large numbers of organoids to the liver without the risk of complications associated with intra-vascular infusion. Concurrently, ectopic locations need to be considered as alternatives to intra-hepatic delivery because this may be inappropriate for certain types of patients with liver disease, such as those with cirrhotic livers, portal hypertension, or significant coagulopathy, which increases the risk of surgical complications [26] and this also creates a hostile environment for transplanted organoids. The vascularised chamber circumvents some of these issues by providing a quick, simple, and relatively minimally invasive surgical technique which creates a closed, protected space in an ectopic location. This surgical device facilitates the reliable generation of substantial volumes of well vascularised tissues and can accommodate large numbers of organoids, unlike other transplantation sites like the spleen or kidney capsule. The chamber can be created around any artery/vein branch, although the groin is commonly used for ease of access [11, 13, 14, 15]. This approach has only been used in a few liver bioengineering studies [12, 13, 14, 15]. The data presented here suggests it has significant advantages over other organoid transplantation methods and its use should be expanded.

Several different mechanisms may influence the survival of organoids at different transplantation sites. This includes the level of vascularisation, hypoxia, immune surveillance, trophic factors, and local microenvironment, as well as whether the organoids were delivered via an injection syringe or pipette, which exposes organoids to different shear forces [27]. For example, reasons for the lack of organoid survival in the spleen could be the relative low levels of extracellular matrix and high immune surveillance in the spleen [28], as well as shear stress exerted on the organoids during injection using a fine needle syringe. Although complex and multi-factorial, these mechanisms need to be explored and optimised to maximise organoid engraftment. One important concept that appears to have considerable impact is constraining organoids within a localised space, using biomaterials. Both the chamber and scaffold achieve this, either in a closed space in the chamber, or within interconnected porous spaces and channels within a scaffold. Interestingly, delivering organoids in a small volume of thermo-responsive hydrogel (Matrigel) alone does not appear to facilitate organoid survival (as seen in the liver, spleen and subcutaneous fat), possibly because the hydrogel is easily dislodged or spread over a large area without physically constraining organoids to the extent that the chamber or scaffold does. Further development of these strategies and investigating the effect of organoid densities, the effect of compacting organoids in a small space, and how this influences viability and overall engraftment is warranted. Additionally, combining the two concepts by transplanting organoids within a scaffold into the chamber may enhance further organoid survival by combining the effects of localising organoids in a small space with the high levels of vascularisation in the chamber. This has been done previously using blood vessels, where the scaffold was used as a carrier vehicle to deliver transplantable blood vessels into the chamber, which acted as a vascularised bioreactor [22].

The type of organoid used for transplantation is also likely to have an important impact on organoid survival and development. There are many varieties of human liver organoids, spanning from the less-clinically relevant immortalised cell-line derived organoids [13, 29] to primary cell-derived organoids [30, 31], and those created from pluripotent stem cells including hiPSC [2, 4]. The cell types within organoids can vary and may be liver progenitor cells [15, 30], cholangiocytes (biliary epithelial cells) [32, 33], hepatocytes [31, 34], or a mixture including non-parenchymal cells such as endothelial or stromal cells [4, 15, 34]. In this study hiPSC-derived organoids were used because these represent the most likely source of cells to be used for future clinical therapy. Furthermore, liver progenitor cells (LPCs) derived from hiPSC were used because they can readily expand in culture unlike terminally differentiated hepatocytes which have a very limited lifespan in culture, and LPCs can form not only hepatocytes but cholangiocytes [30, 35] which are important for the drainage of bile produced by hepatocytes. In our system, the chamber provides the most optimal environment for iLPC organoids, with limited to no survival in the spleen and subcutaneous group. However, in other studies, different types of liver organoids have been reported to survive after intra-splenic injection by translocating into the liver [3], and in subcutaneous fat [36] (although not compared to other sites). These various types of organoids should be tested in the chamber device. Furthermore, when considering other organoid transplantation studies, it is worth noting that many organoid transplantation studies do not transplant whole organoids, but organoid-derived single cells [30, 32, 37, 38]. This effectively serves as cell therapy, not organoid therapy, and makes it difficult to compare these studies to those transplanting actual organoids.

One interesting aspect of this study worth further developing is the use of the polyurethane scaffold loaded with iLPC organoids, which can serve as a “liver patch” that can be applied on the liver surface. Several other studies have reported liver cell/organoid patches [39, 40, 41, 42, 43], and the advantage of such methods is the ability to easily apply this using minimally invasive methods such as laparoscopic surgery, and this can be delivered on multiple areas of the liver surface, and even repeated over time. Most of these studies have used extra-cellular matrix and synthetic hydrogels [39, 40], stacked cell sheets [43] or electro-spun fibres [42] to create the patches, however these methods create thin delicate patches that can be difficult to physically manipulate without damaging them. The advantage of the polyurethane scaffold is its robust physical properties. The scaffold is a sponge-like foam that can be easily picked up, loaded with organoids, and transferred to the liver surface in one piece, without any risk of tearing or disintegrating. This means it can be upscaled in diameter and thickness and still be easily applied to the liver surface and will physically hold its form. Although an attractive concept, given that organoid survival in these scaffolds was relatively poor in this study, there needs to be further optimisation to increase organoid survival. This may involve increasing the organoid number seeded per scaffold and incorporating additional non-parenchymal cells that enhance organoid survival and maturation [15, 44, 45].

Future directions for this work include assessing longer time-points to determine the extended survival of organoids with the vascularised chamber and intra-hepatic scaffold, repeating the study with different kinds of liver organoids, and comparing transplantation sites across different animal models to confirm these findings are universally applicable. An additional transplantation site that should be considered is the omentum, which has been often used to transplant organoids and hepatocytes including in human studies [7, 8, 34]. It is an easily accessible large space which is well vascularised and directly connected to the portal blood supply. However, one major limitation of this approach is that any major intra-abdominal surgery predisposes the patient to intra-abdominal adhesions (scarring), which often causes long-term complications such as repeated bowel obstructions [46]. Furthermore, any infection or tumour formation can lead to rapid peritoneal dissemination. It is also likely that organoids will need to be delivered using a scaffold, otherwise they can be easily dislodged without attaching to any surface. These issues need to be carefully considered in future studies. Lastly, the most effective methods of organoid transplantation need to be applied in different models of liver disease to stratify which approach is best for a type of liver disease.

Overall, this study adds clinically relevant data to the field of liver organoid transplantation by highlighting modalities of organoid transplantation that optimise organoid survival and suggesting a roadmap for future developments to hasten the clinical translation of this concept.

## ACKNOWLEDGEMENTS

The authors thank Anna Deftereos and Amanda Rixon from the Experimental Medicine and Surgery Unit, St Vincent’s Hospital Melbourne for assistance with animal care and experiments, and Dr Chandana Herath (University of Melbourne), Professor Ian Alexander (University of Sydney) and Yecuris Corporation for providing FRG mice. The authors also acknowledge the Monash Organoid Program for providing Wnt3A, R-Spondin1 and Noggin conditioned medium, and Professor Eduoard Stanley and Professor Andrew Elefanty (Murdoch Children’s Research Institute) for providing the eGFP hiPSC line. The purified GCTM-5 antibody used to purify iLPCs was produced using a hybridoma line from Professor Martin Pera (Jackson Laboratory) and Dr Alison Farley (Walter and Eliza Hall Institute). The polyurethane scaffold (NovoSorb®) used in the study was provided by PolyNovo Ltd.

The authors received funding from the Australian National Health & Medical Research Council (NHMRC), St Vincent’s Hospital Melbourne Research Endowment Fund, Stafford Fox Medical Research Foundation, O’Brien Foundation, St Vincent’s Institute Foundation, Australian and New Zealand Hepatic, Pancreatic and Biliary Association, University of Melbourne Centre for Stem Cell Systems, Bioplatforms Australia, and the Victorian State Government’s Department of Business Innovation Operational Infrastructure Support Program.

## Notes

### Competing Interest Statement

The authors have declared no competing interest.

